# Phenotypic and environmental predictors of reproductive success in painted turtles

**DOI:** 10.1101/2020.11.10.377028

**Authors:** Jessica M. Judson, Luke A. Hoekstra, Kaitlyn G. Holden, Fredric J. Janzen

## Abstract

Sexual selection is often assumed to elicit sexually dimorphic traits. However, most work on this assumption in tetrapod vertebrates has focused on birds. In this field experiment, we assessed relationships between both sexually dimorphic (body size, claw length) and non-dimorphic traits (forelimb stripe color, baseline corticosterone concentrations) and reproductive success in adult painted turtles to explicate the roles of these phenotypes in mate choice and the evolution of sexual dimorphism. We also modified adult sex ratios in experimental ponds to elucidate the role of biased sex ratios on reproductive success, which is a timely test of the potential threat of biased sex ratios on population persistence in a species with temperature-dependent sex determination. We found no strong influence of male phenotypes on male siring success, but female body size and baseline corticosterone concentrations predicted female clutch sizes. We find weak evidence that adult sex ratio influences male siring success, with a male-biased sex ratio producing lower male siring success than a female-biased sex ratio. This study offers evidence that female mate choice may not be an important selective force on male phenotypes, but that instead selection occurs on female phenotypes, particularly body size and corticosterone concentrations. Further, biased adult sex ratios can influence reproductive success of both sexes. Finally, the use of Kompetitive Allele Specific PCR (KASP) was highly successful in parentage analysis, which adds reptiles to the growing list of taxa successfully genotyped with this new technology.

**Lay Summary:** Female painted turtles aren’t choosy about traits of their mates. In a field experiment, we find that male traits do not predict male fitness, but key female traits (body size and stress levels) do predict female reproductive success. Further, we find weak evidence that adult sex ratio influences individual fitness in this species with environmental sex determination. Ultimately, we reject the long-assumed importance of female mate choice in this freshwater turtle.

## Introduction

Sexually dimorphic traits frequently inspire studies of sexual selection. While some sexually dimorphic traits can be explained by sexual selection on males (e.g., White et al. 2018), others do not influence male reproductive success, and instead may reflect selection on females (Blanckenhorn 2005). Phenotypes that do not exhibit sexual dimorphism can also influence male or female reproductive success (e.g., Kelly et al. 2012). Furthermore, the environmental context of mating, such as adult sex ratio, can affect reproductive success, shaping population dynamics and persistence (Szekely et al. 2014).

Large body size is typically a strong predictor of male reproductive success (e.g., Shine et al. 2000; White et al. 2018), particularly in species where male-male competition for mates or territories occurs, or where forced matings are common. However, under female-biased sexual size dimorphism (SSD), female choice may determine reproductive success (Berry and Shine 1980). Male body size may still be important for mate choice in such species, but female-biased SSD can evolve due to fecundity selection on females (Blanckenhorn 2005). In these situations, large female body size predicts increased reproductive output (e.g., Cox et al. 2003). In addition to SSD, other sexually dimorphic traits, including color, can affect reproductive success. Male color influences reproductive success in many vertebrates (e.g., Siefferman and Hill 2003; Salvador et al. 2007).

Non-dimorphic phenotypes also may be important for reproductive success of males and females. In species with high biparental investment in care of offspring, both sexes should be choosy in selecting mates (Johnstone et al. 1996). Brightly colored ornaments exhibited by both sexes can send similar or different signals of quality to potential mates (Kelly et al. 2012). However, in vertebrates without heavy investment in parental care, the relationship between color and reproductive success is less well understood. In some species of brightly colored pond turtles, for example, color may not vary between the sexes (Judson et al. In review), yet female choice of males is often invoked as generating bright colors on the skin and plastron (e.g., Polo-Cavia et al. 2013). Thus, color may still play a role in reproductive success even when color is not sexually dimorphic.

Beyond morphology, physiological phenotypes associated with stress responses may predict reproductive success. Stress hormones (i.e., glucocorticoids) in vertebrates are essential mediators of energy balance in response to both acute stressors and other common activities, including feeding and reproduction (Landys et al. 2006). Glucocorticoids are often studied in the context of trade-offs, as acutely or chronically stressed iteroparous organisms may need to allocate energetic resources toward survival at the expense of reproduction (Wingfield and Sapolsky 2003). The CORT-Fitness Hypothesis (Bonier et al. 2009) posits that stressed vertebrates may experience decreased fitness as a result of increased stress (often measured by concentrations of corticosterone; CORT). However, increased CORT during reproductive activity can also facilitate reproductive behaviors and have positive effects on fitness (Bonier et al. 2009). Thus, elevation of glucocorticoids is not in and of itself indicative of a negative acute or chronic life event, and context matters. Although CORT can directly mediate reproductive physiology and behavior, it can also interact with mating signals, including color, to influence mate choice and, ultimately, fitness (Moore and Hopkins 2009; reviewed in Leary and Baugh 2020).

In addition to phenotype, environmental conditions can substantially influence individual reproductive success. Resource availability often strongly determines reproductive success (e.g., Hoset et al. 2017). In some populations, the adult sex ratio (ASR) may be biased, thus the availability of mates may be an important resource dictating reproductive success and future population dynamics. Biased ASR can modify the frequency of intrasexual (Weir et al. 2011) and intersexual competition (Le Galliard et al. 2005) and change the dynamics of mate choice (Atwell and Wagner 2014; Grant and Grant 2019), which can have long-term consequences for population persistence (e.g., Steifetten and Dale 2006; reviewed in Szekely et al. 2014). In common lizards, for example, increased competition due to skewed ASR changed behavior and increased intersexual aggression, leading to population declines (Le Galliard et al. 2005). Many vulnerable species exhibit biased ASR due to skewed death rates of males or females (e.g., Heinsohn et al. 2019). In reptiles with temperature-dependent sex determination (TSD), biased ASR might be exacerbated by ongoing climate warming producing skewed offspring sex ratios (Janzen 1994; Schwanz et al. 2010). These biased ASRs also could contribute to decreased effective population sizes and trap populations in an extinction vortex (Grayson et al. 2014). Understanding the influence of skewed ASR on reproductive success, particularly in species with TSD, is thus important in both basic and applied contexts.

In this study of a pond turtle with female-biased SSD, we measured body size, claw length, forelimb stripe color, and baseline CORT concentrations to quantify their relative influence on male and female reproductive success. Furthermore, using semi-natural experimental ponds, we modified the ASR to test its influence on reproductive success and potential impact on population dynamics of a species with TSD. Finally, we assigned parentage of offspring by developing a set of SNPs from population-level RADseq data and used a genotyping technology new to reptiles, Kompetitive Allele Specific PCR (KASP), to genotype all individuals included in this study.

## Methods

### Study System

The painted turtle (*Chrysemys picta*) is widespread in North America (Ernst and Lovich 2009). Adults are sexually dimorphic: males have elongated foreclaws and females are larger, suggesting forcible insemination is likely uncommon (Berry and Shine 1980; but see Hawkshaw et al. 2019). Visual and tactile courtship displays performed before mating offer the potential for female choice based on male traits, including his body size, claw length, and color (Ernst and Lovich 2009). Females can store sperm, which could allow for cryptic female choice, and some clutches in the wild exhibit multiple paternity (~13%, Pearse et al. 2001; >30%, Pearse et al. 2002; ~14.1%, McGuire et al. 2014), typically with two sires represented in a single clutch. Males may choose females based upon female size (McGuire et al. 2014), as size is an indicator of female reproductive output (e.g., Hoekstra et al. 2018), or by coloration (Judson et al. In review). Alternatively, males may instead attempt mating with as many females as possible (Bateman 1948) based on encounter rate, or may mate randomly with respect to female traits. Predictors of reproductive success for painted turtles beyond female size remain elusive. The brightness and hue of adult forelimb stripes indicate aspects of stress and immune health (Judson et al. In review), and thus could signal mate quality, though the influence of color on reproductive success has not been evaluated. Moreover, social context, especially the ASR (e.g., Szekely et al. 2014), can influence mating systems generally, but its influence on reproductive success in painted turtles is not well understood. Incubation temperature determines hatchling sex ratios in painted turtles (e.g., Janzen 1994), which in turn affect ASR (Schwanz et al. 2010), and climate change models predict a warming environment that could further skew sex ratios over time (Refsnider and Janzen 2016).

### Turtle Husbandry and Sampling

The following research methods were approved by Iowa State University (ISU) IACUC (12-03-5570-J). We constructed three semi-natural experimental ponds (19m × 15m × 1.5m) at the ISU Horticulture Farm surrounded by 25m x 55m x 1m silt fencing with aluminum flashing to prevent movement of turtles between ponds and to allow females adequate area surrounding the ponds to construct nests (Judson et al. In review; Fig. 1). In April 2016, we released 63 adults (26 females, 37 males), collected from the Thomson Causeway Recreation Area (TCRA) in Thomson, IL, USA, into the ponds. Permits for painted turtle collection were provided by the United States Army Corps of Engineers, the United States Fish and Wildlife Service (SUP 32576-021), and the Illinois Department of Natural Resources (NH11.0073). We classified all turtles as sexually mature using a combination of size, presence of sexually-dimorphic characters (e.g. elongated claws in males), and annual rings on plastral scutes (Moll 1973). We placed turtles into the ponds with differing ecologically-relevant ratios of females to males recorded in wild populations (e.g., Hughes 2011; Dupuis-Désormeaux et al. 2017), which we refer to as an ASR treatment: male-biased (M > F), female-biased (M < F), and approximately equal number of males and females (M = F; Fig. 1). We supplemented the turtles’ diet with Mazuri^®^ Aquatic Turtle Diet for the duration of the experiment, which did not measurably change forelimb stripe coloration between captive and wild turtles (Judson et al. In review). In May and June of 2016, we monitored ponds for nesting activity hourly 0600-2200 h. After a female finished nesting, we obtained a blood sample with a heparin-rinsed syringe from the caudal vein for genotyping. We removed eggs from each nest, placed them into plastic containers filled with moist vermiculite (−150 kPa), and incubated them at 28°C until hatching at ISU. Following incubation, we sampled tissue from 207 hatchlings for genotyping, excluding undeveloped embryos and infertile eggs (N=20).

**Figure 1:**
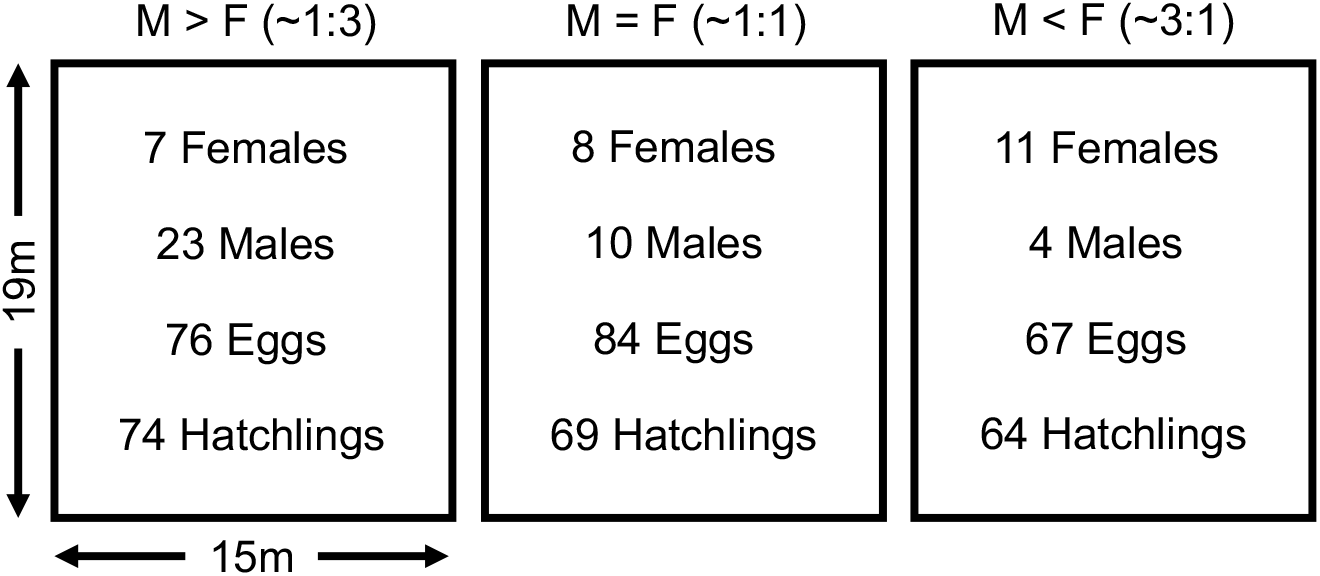
Diagram of adult sex ratio treatment. Number of female and male painted turtles released into each pond, the number of eggs laid, and number of offspring successfully hatched from each pond. Arrows indicate dimensions of each pond in meters.

In July 2016, following the nesting season, we removed the turtles from the ponds and collected a blood sample within 10 minutes of capture. This 10-minute limit was to assess baseline measures of CORT in the bloodstream, which rise after 10 minutes of handling in painted turtles (Polich 2016). We centrifuged the blood to separate the plasma before snap freezing plasma and red blood cells in liquid nitrogen and storing at −80°C for CORT measures and genotyping, respectively. We also measured other aspects of stress and immune function (Judson et al. In review), but we do not consider these variables here, as we did not have strong a priori predictions for their influence on reproductive output in painted turtles, and post-hoc analysis of the effects of unreported physiological variables on reproductive success did not affect our conclusions regarding the effects of CORT. Finally, we measured plastron length for all turtles and length of the third claw of each forelimb to the nearest mm for males, which is usually the longest claw (McTaggart 2000; Hughes 2011). We averaged the two claw measures, excluding any claws that were broken in our averages (N=5).

### Color Analysis

For color analysis, we followed the methods of Judson et al. (In review). Briefly, we used a tripod-mounted Canon EOS Digital Rebel XSi camera and EF-S18-55mm lens to take RAW-formatted photographs of each turtle’s cranial region under controlled incandescent lighting with a grey standard (18% reflectance; Insignia NS-DWB3M) in every photograph. We used the Image Calibration and Analysis Toolbox v. 1.22 (Troscianko and Stevens 2015) in ImageJ v. 1.52a (Schneider et al. 2012) to linearize photographs and obtain reflectance measures. We measured reflectance as close as possible to the middle point of the right forelimb stripe, which appears yellow, orange, or red to the human eye. We performed the above process for all except six turtles, who either were ill (N=2: 1 female, 1 male) and thus not photographed, or were not recovered from the ponds (N=4 females, likely due to predation; F. Janzen personal observation) and thus were not photographed or measured for CORT. We used two measures to represent color variation of forelimb stripes with long wavelength (LW), medium wavelength (MW) and short wavelength (SW) reflectance measures: overall percent brightness, calculated as 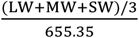 (Troscianko and Stevens 2015; Judson et al. In review), and hue, calculated as LW/MW (Judson et al. In review). High values of percent brightness indicate lighter stripe color, whereas high values of hue indicate increased redness and decreased yellowness of the forelimb stripe.

### Corticosterone

CORT levels reflect vertebrate stress responses, but also function in concert with other physiological factors to facilitate reproductive behaviors, feeding, and maintenance of homeostasis (Moore and Jessop 2003; Landys et al. 2006). To quantify baseline concentrations of circulating plasma CORT (ng/mL), we used a double-antibody radioimmunoassay (ImmuChem Double Antibody Corticosterone I-125 RIA kit, MP Biomedicals, Irvine, CA, USA (Polich 2016; Judson et al. In review). We quantified samples (N=59) in duplicate and included a pooled sample to assess inter-assay variability (average coefficient of variation 4.0%).

### Parentage Analysis

We used Kompetitive Allele Specific PCR (KASP) all-inclusive services to extract DNA and genotype 96 SNP loci in all 63 adults and 207 hatchlings (Semagn et al. 2014). KASP has been successful for genotyping many crop species (McCouch et al. 2010; Khera et al. 2013), and its use in vertebrates is increasingly common (e.g., Wielstra et al. 2016; Bourgeois et al. 2018). To determine the 96 SNP loci that confidently assign parentage, we used RADseq data of known parent-offspring pairs from this population (FJ Janzen, unpublished data). Our SNP filtering of RADseq data was adapted from GATK Best Practices (McKenna et al. 2010; DePristo et al. 2011). Briefly, we used a minGQ filter of 20, kept only biallelic sites, allowed only 1% missing genotypes for each site, and used a minor allele frequency filter of 0.4 to select SNPs with high heterozygosity in the population. This filtering yielded 801 SNPs. Next, we used the SAMtools v. 1.4 (Li et al. 2009) faidx command to query the surrounding sequence of each SNP from the *C. picta* draft genome v. 3.0.3 (Shaffer et al. 2013). We removed any SNPs with missing data or ambiguous sequence in the 50 bp upstream or downstream of the SNP of interest. We further reduced the SNP set by removing SNPs of interest that had >1 SNP in the flanking regions for a final set of N=150. We measured linkage disequilibrium of these SNPs in PLINK v. 1.9 (Purcell et al. 2007) to ensure that none were linked (*r*^2^ < 0.5, average *r*^2^ across all SNPs = 0.04). We selected 96 SNPs from this set, a number that yielded high parentage assignment success in other studies of vertebrates (Hauser et al. 2011). We compared flanking primer sequences for these SNPs to the *C. picta* draft genome using BLAST to eliminate multiple matches.

To analyze genotypes obtained from KASP, we used the pedigree program Cervus v. 3.0.7 (Kalinowski et al. 2007). We recorded maternity assignment for all clutches during nesting observations, and Cervus assigned these recorded mothers to the correct clutches in all cases. Thus, we tested paternity with known maternal genotypes in Cervus with default settings. We also analyzed sibship among hatchlings using the full likelihood method of COLONY v. 2.0.6.5 (Jones and Wang 2010) to confirm multiple paternity and provide insight into relationships among hatchlings resulting from sperm storage that were not sired by males included in the experimental ponds.

### Statistics

For all statistical analyses and plotting, we used R version 4.0.2 (R Core Team 2020). Our final sample sizes for inclusion in statistical analyses were 22 females and 37 males recovered from the ponds (see Color Analysis). As we expected that predictors of reproductive success would differ between males and females, we modeled the sexes separately. We standardized the continuous predictor variables (plastron length, CORT concentrations, forelimb stripe brightness and hue, and male claw length) by sex to mean of zero and unit variance so that slopes could be directly compared among variables (Grueber et al. 2011). We checked for outliers in continuous predictors using a threshold of three standard deviations from the mean for each sex, and found one male outlier for forelimb stripe brightness, which we removed.

Female reproductive success - We assessed correlations among female measures, and found that no predictor variables were strongly correlated (−0.63 < *r* < 0.52). For females, the measure of reproductive success was the clutch size (including infertile eggs and undeveloped embryos), and the full model conditional upon female clutch size being greater than zero was as follows:

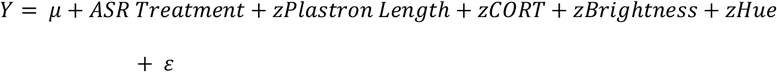

where *μ* represents the grand mean and *ε* the error term, and “z” precedes standardized continuous predictors. We included ASR treatment as a fixed effect. We included plastron length, as body size is an important predictor of clutch size in female painted turtles (e.g., Hoekstra et al. 2018). Clutch size was under-dispersed, as is typical of reproductive data (Brooks et al. 2019), and three females that were recovered from the ponds did not oviposit. Thus, to account for a zero-inflated, under-dispersed count distribution, we assessed generalized linear models using ‘glmmTMB’ v 1.0.2.1 (Brooks et al. 2017; Brooks et al. 2019). We used an all-subset approach to model selection, which included every combination of variables from the full model and intercept-only models which assess only the constant and residual variance (Grueber et al. 2011), with a zero-inflated Conway-Maxwell-Poisson error distribution to account for under-dispersion and zero-inflation in clutch size using ‘dredge’ from ‘MuMIn’ v 1.43.17 (Bartoń 2020). The zero-inflation model included an intercept-only zero-inflation model (~1; Brooks et al. 2019) with no other predictors included, as only a small number of females did not reproduce, and increased zero-inflation model complexity induced model convergence issues. We selected the best-fitting model(s) using AICc (Burnham and Anderson 2002). To compare models using AICc, we removed the female for which we did not have color measures from all model comparisons, as comparisons among models are only valid when the same data are included for each model (Symonds and Moussalli 2011). We tested the full and best-fitting models for dispersion and model fit with ‘DHARMa’ v 0.3.3.0 (Hartig 2020) and for multicollinearity using ‘performance’ v 0.5.0 (Lüdecke et al. 2020). We also calculated Akaike model and parameter weights for all model subsets using ‘MuMIn’ v 1.43.17. We report parameter estimates for the models within two ΔAIC_c_ units from the best submodel, excluding any nested models included in that threshold (Arnold 2010).

Male reproductive success - For males, we checked correlations and assessed multicollinearity in the same manner as for females. We found no strong correlations amongst predictor variables (−0.58 < *r* < 0.53). The measure of reproductive success was the number of offspring sired (i.e., absolute fitness; e.g. Noble et al. 2013) as determined by parentage analysis, and the full model conditional upon the number of offspring sired being greater than zero was as follows:

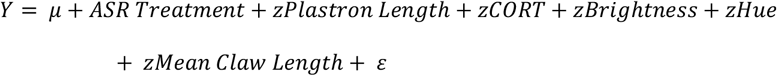

We modeled males in the same manner as females with a few exceptions. Number of offspring sired was both zero-inflated and over-dispersed, thus we assessed full conditional model fit with zero-inflated Poisson, Conway-Maxwell-Poisson, and negative binomial error distributions. A zero-inflated Conway-Maxwell-Poisson error distribution best fit the full conditional model according to dispersion tests in ‘DHARMa’ v 0.3.3.0 and comparison among error distributions using AICc, so we used this error distribution for all model subsetting comparisons. We excluded three males due to missing data from subsetted models (see Color Analysis). The zero-inflation model for males included the same predictors as the male full conditional model, and intercept-only conditional and zero-inflation models were included in model comparisons. We also performed model subsetting comparisons with an intercept-only zero-inflation model for comparison. We created all figures of results using ‘ggplot2’ v. 3.3.2 (Wickham 2016).

Opportunity for Selection - Finally, we measured the Opportunity for Selection using a new index, Δ_*I*_, which allows comparison between males and females when sex ratios are unequal (Waples 2020). We used the same offspring life stage comparison for males and females, the number of hatchlings, for this index.

## Results

The 26 females laid 22 clutches during the nesting season, resulting in 227 eggs laid. Clutch sizes ranged from 0-14 eggs, with a mean of 10.3 eggs among ovipositing females (Fig. 2). Of the 227 eggs laid, 207 offspring successfully hatched, split between the male-biased pond (74 offspring), equal sex-ratio pond (69 offspring), and female-biased pond (64 offspring; Fig. 1).

**Figure 2:**
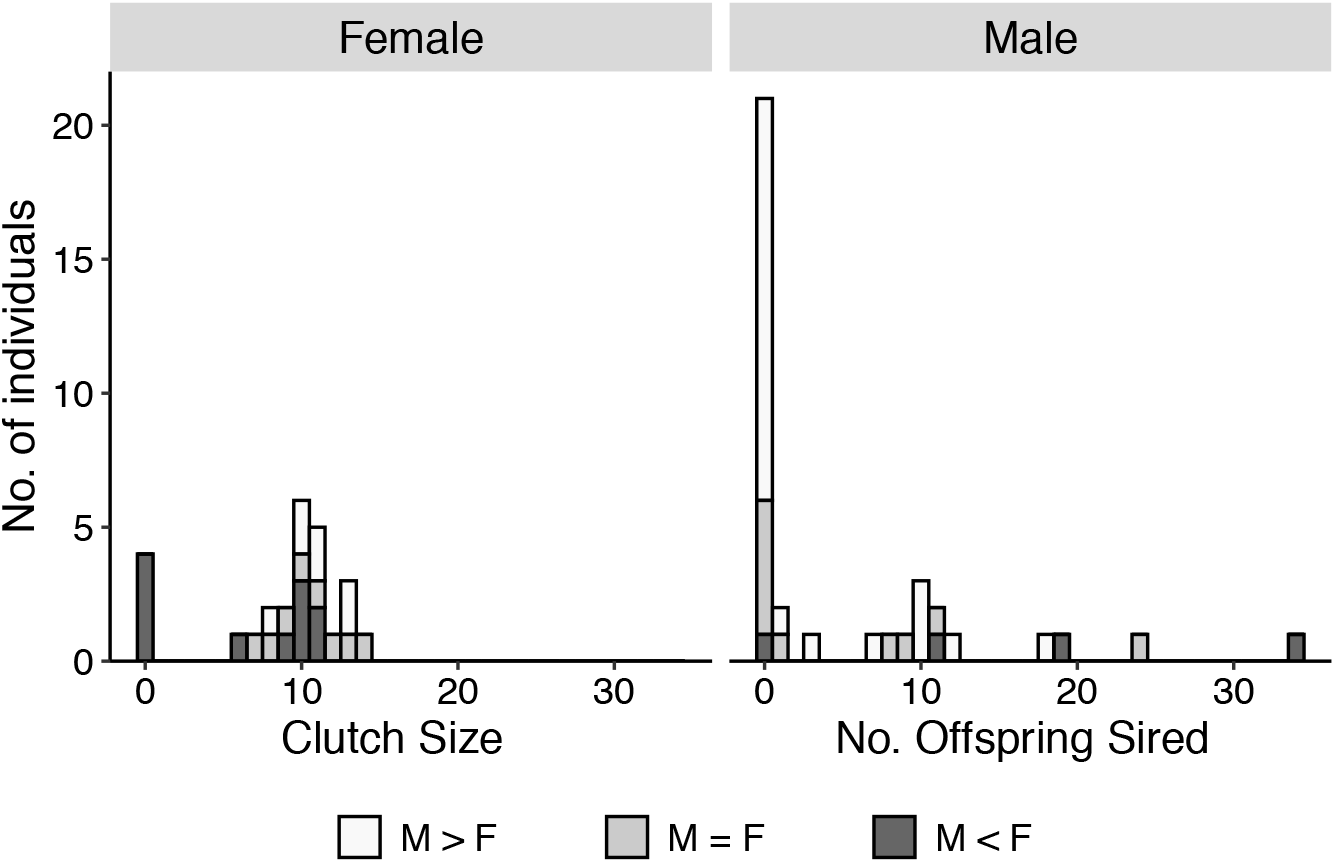
Stacked histogram of female and male reproductive output for adult painted turtles in this study (N=63) labeled by adult sex ratio treatment.

The final panel of SNPs consisted of 88 loci, as 8 of the 96 loci were not successfully genotyped due to unclear separation of clusters or poor amplification during KASP genotyping. We genotyped all 207 hatchlings, though 5 individuals genotyped at fewer than 50 loci could not be assigned to a single sire. These 5 hatchlings were thus excluded from statistical analyses of males. We assigned paternity for all remaining hatchlings with high confidence, with the exception of two clutches (14 hatchlings) that did not have likely sires among the males in this study. Offspring within each of these two clutches were full siblings with a probability of 1; there was no evidence of half siblings between the two clutches, suggesting that the sires were different individuals whose sperm was stored and that these clutches did not exhibit multiple paternity. Among the males in our study, 21 sired no offspring, and 16 sired between 1 and 34 offspring in 1 to 4 different clutches, with a mean of 11.8 offspring sired (Fig. 2). The incidence of multiple paternity was 18%, or 4 out of 22 clutches, with two multiply-sired clutches each from the male-biased and equal-ratio ponds. Opportunity for Selection (Δ_*I*_) was 0.12 for females and 2.33 for males.

### Females

The best-fitting model according to AICc included plastron length and CORT (Table 1, Table S1). Larger females and those with lower CORT concentrations laid more eggs (Table 2; Fig. 3). Pond ASR and females’ forelimb stripe brightness and hue did not predict female reproductive success, as indicated by low Akaike weights of these variables (0.09, 0.14, and 0.24, respectively) compared to plastron length (0.99) and CORT (0.46). We detected no multicollinearity in either the full or best-fitting model nor issues with model fit, as dispersion tests were non-significant (Figs. S1, S2).

**Figure 3:**
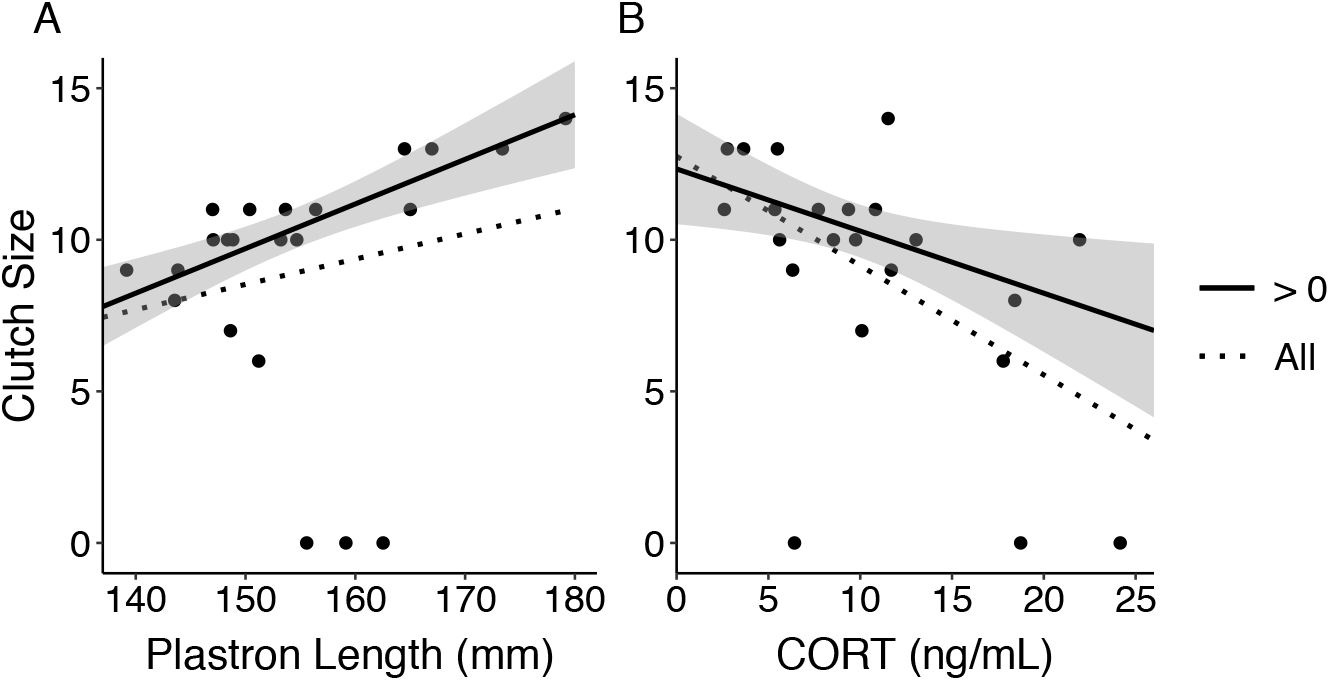
Relationship between number of eggs laid by female painted turtles and plastron length (A) or corticosterone concentrations (CORT; B). Raw values are plotted, and lines depict a simple linear regression using all females (All, dashed line) or excluding females that did not lay eggs (> 0, solid line). Gray shading depicts 95% confidence interval.

**Table 1:** Full, intercept-only, and 5 best-fitting models for female and male painted turtle reproductive success ranked according to AIC_c_

| Conditional Model | Zero-inflation Model | df | ΔAIC<sub>c</sub> | Weight |
| --- | --- | --- | --- | --- |
| **Females** |  |  |  |  |
| ~ PL<sup>1</sup> + CORT<sup>2</sup> | ~ 1 | 5 | 0.00 | 0.33 |
| ~ PL | ~ 1 | 4 | 0.41 | 0.27 |
| ~ PL + Hue | ~ 1 | 5 | 1.86 | 0.13 |
| ~ PL + CORT + Brightness | ~ 1 | 6 | 3.75 | 0.05 |
| ~ PL + CORT + Hue | ~ 1 | 6 | 3.76 | 0.05 |
| ~ 1 | ~ 1 | 3 | 12.20 | 0.00 |
| ~ ASR<sup>3</sup> + PL + CORT + Brightness + Hue | ~ 1 | 9 | 15.73 | 0.00 |
| **Males** |  |  |  |  |
| ~ ASR | ~ 1 | 5 | 0.00 | 0.05 |
| ~ Brightness | ~ 1 | 4 | 0.39 | 0.04 |
| ~ ASR + Brightness | ~ 1 | 6 | 1.70 | 0.02 |
| ~ ASR + Hue | ~ 1 | 6 | 1.82 | 0.02 |
| ~ Brightness + CORT | ~ 1 | 5 | 1.96 | 0.02 |
| ~ 1 | ~ 1 | 3 | 2.50 | 0.01 |
| ~ ASR + PL + CORT + Brightness + Hue + MCL<sup>4</sup> | ~ ASR + PL + CORT + Brightness + Hue + MCL | 17 | 53.43 | 0.00 |

All continuous predictors standardized by sex. <sup>1</sup>plastron length; <sup>2</sup>baseline corticosterone concentration; <sup>3</sup>adult sex ratio treatment; <sup>4</sup>mean claw length

**Table 2:** Parameter estimates for models of female painted turtle reproductive success within two ΔAIC_c_ units of the best-fitting submodel

| Parameters | Estimate | SE | z |
| --- | --- | --- | --- |
| **Model 1 ($\Delta AIC_c=0$)** |  |  |  |
| **<u>Conditional Model</u>** |  |  |  |
| Intercept | 2.32 | 0.03 | 80.92 |
| PL<sup>1</sup> | 0.12 | 0.03 | 4.27 |
| CORT<sup>2</sup> | -0.07 | 0.04 | -2.07 |
| **<u>Zero-inflation Model</u>** |  |  |  |
| Intercept | -2.25 | 0.74 | -3.03 |
| **Model 2 ($\Delta AIC_c=0.41$)** |  |  |  |
| **<u>Conditional Model</u>** |  |  |  |
| Intercept | 2.34 | 0.03 | 76.39 |
| PL | 0.13 | 0.03 | 4.83 |
| **<u>Zero-inflation Model</u>** |  |  |  |
| Intercept | -2.25 | 0.74 | -3.03 |

### Males

Best-fitting models according to AICc included either ASR treatment or forelimb stripe brightness (Table 1; Table S2). However, model weights were low for all models (≤ 0.05), and parameter weights were all ≤ 0.50, suggesting considerable model uncertainty (Table S2). Additionally, the null model including only intercepts for the conditional and zero-inflation parameter was 2.5 ΔAIC_c_ units from the best-fitting model (Table 1), suggesting the included variables are weak predictors of male reproductive success. Notwithstanding, according to these models, female-biased ASR and greater forelimb stripe brightness increased individual reproductive success (Table 3, Fig. 4). As with the female models, we detected no multicollinearity in conditional and zero-inflated full and best-fitting models and no issues with model fit as indicated by non-significant dispersion tests (Figs. S3-S4). Model ΔAIC_c_ rankings for the best-fitting models with an intercept-only zero-inflation model used in model subsetting were the same as those including a zero-inflation model that matched the full conditional model (Table S3).

**Figure 4:**
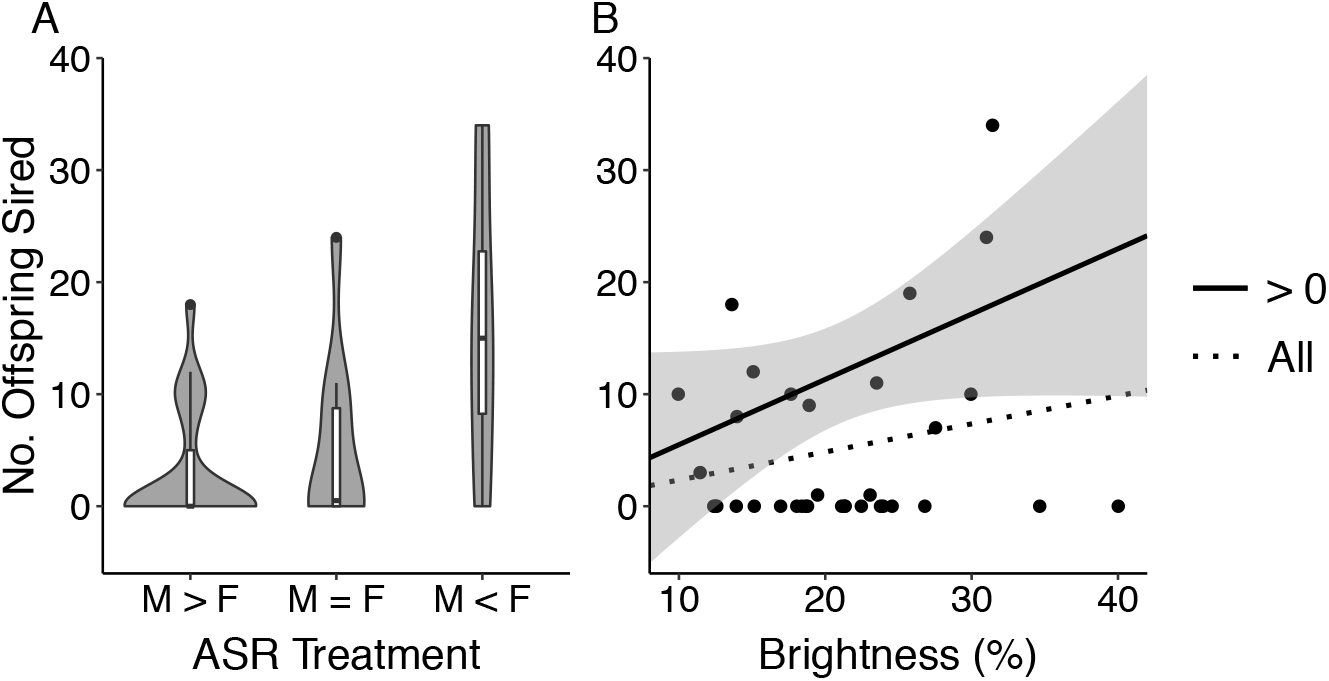
Violin plot for number of offspring sired by male painted turtles from each adult sex ratio (ASR) treatment (A), and relationship between male percent brightness of the forelimb stripe and number of offspring sired (B). Boxplots are inset within violin plots, and raw values are plotted for both panels. Lines in panel B depict a simple linear regression using all males (All) or excluding males that did not sire offspring (> 0). Gray shading depicts 95% confidence interval. One outlier for brightness was removed.

**Table 3:** Parameter estimates for models of male painted turtle reproductive success within two ΔAIC_c_ units of the best-fitting submodel

| Parameters | Estimate | SE | z |
| --- | --- | --- | --- |
| **Model 1 ($\Delta AIC_c=0$)** |  |  |  |
| **<u>Conditional Model</u>** |  |  |  |
| Intercept | 2.56 | 0.25 | 10.41 |
| ASR<sup>1</sup>: F > M | 0.72 | 0.35 | 2.04 |
| ASR: M > F | -0.40 | 0.32 | -1.24 |
| **<u>Zero-inflation Model</u>** |  |  |  |
| Intercept | 0.33 | 0.35 | 0.95 |
| **Model 2 ($\Delta AIC_c=0.39$)** |  |  |  |
| **<u>Conditional Model</u>** |  |  |  |
| Intercept | 2.51 | 0.15 | 16.35 |
| Brightness | 0.42 | 0.18 | 2.40 |
| **<u>Zero-inflation Model</u>** |  |  |  |
| Intercept | 0.33 | 0.35 | 0.93 |

More complex submodels excluded when nested submodels had lower AIC values; <sup>1</sup>adult sex ratio treatment

## Discussion

We quantified two sexually-dimorphic traits, body size and male claw length, in addition to plasma CORT and color of forelimb stripes in adult painted turtles in a field experiment in which we manipulated ASR to assess the impact of these factors on measures of individual fitness. Female reproductive success was positively related to body size and negatively related to plasma CORT, whereas male reproductive success was not strongly predicted by any measured phenotypes or by ASR. Our results suggest directional selection on morphology and physiology in females, and are thus congruent with the perspective that sexual dimorphism in traits could arise from selection on just one sex, rather than on both (Janzen and Paukstis 1991).

The ratio of reproducing males to females was approximately 1:1.4, with 22 of 26 females and 16 of 37 males successfully reproducing. Reproductive success varied widely among turtles, but was more skewed in males than females, such that the Opportunity for Selection (Δ_*I*_) was much greater in males than females. Most females reproduced, and the mean number of eggs laid by nesting females (10.4) and the range of eggs laid (0-14) were similar to results of other studies of painted turtles from the TCRA population (mean 10.9 eggs, range 1-14 per nest, Pearse et al. 2002; mean ~10 eggs per nest across lifetime, Delaney et al. In Press). For males, reproductive success ranged from 0 to 34 offspring, with a mean of 11.8 offspring sired across successful males, and fewer than half of the males in this study sired any offspring. These results are similar to a 4-year study of painted turtles from southeastern Michigan, which found that successful males sired on average 8.6 offspring with a range of 1 to 32 offspring (McGuire et al. 2014). Given the considerable variation in male reproductive success, our experiment had ample scope to identify any selection on the male traits we measured, yet we found none.

We also detected multiple paternity in 18% of the clutches, which is similar to estimates of multiple paternity prevalence in free-ranging painted turtles from this population (10.7% observed, 30.1% estimated, Pearse et al. 2002). Interestingly, despite this experiment taking place following the third summer of holding turtles in the ASR treatment ponds (Judson et al. In review) to prevent the use of stored sperm, we found two instances of probable long-term sperm storage in the equal ASR pond, as the genotypes of two sires were not matched by the males in our experiment. Sperm storage has been documented for up to three years in female painted turtles at the TCRA, and recent matings sire the initial clutches of offspring in a ‘last in, first out’ pattern (Pearse et al. 2001). Thus, the two females in our study probably did not mate with males in their pond, and instead utilized stored sperm. The reason for this is unclear, but the decision to use stored sperm deserves further study in wild populations. Overall, given the similar patterns of reproductive success in our experiment and studies of wild populations, our husbandry of turtles should be representative of potential phenotypes influencing reproductive success in the wild.

Sexually-dimorphic phenotypes - We investigated two sexually dimorphic phenotypes in this study, body size (as measured by plastron length) and male claw length, to assess their role in reproductive success. Body size was strongly positively associated with the number of eggs laid by females. This finding is consistent with prior studies (e.g., McGuire et al. 2014; Hoekstra et al. 2018) and aligns with phylogenetic analyses supporting the role of fecundity selection in the evolution of SSD in emydid turtles (e.g., Stephens and Wiens 2009). In contrast, male body size and male claw length did not predict male siring success, and body size was not predictive of whether a male sired offspring or not (t-test *P*=0.79). Similarly, carapace length (which is strongly correlated with plastron length, Hoekstra et al. 2018) did not differ between successful and unsuccessful free-ranging males at the TCRA in an earlier study (Pearse et al. 2002). As plastron length is a proxy for age in this population (Hoekstra et al. 2018), these results imply that female reproductive success increases with age due to increasing body size, whereas male reproductive success does not increase with age. Successful male painted turtles from northwestern Ontario had shorter carapaces (McTaggart 2000), but another study of the same population found no clear relationship between male plastron length and reproductive success (Hughes 2011). Thus, small males appear to accrue no reproductive advantage, and female choice of male body size is not supported in this study. If male coercion were important for copulation, large males would be expected to have increased reproductive success (Hawkshaw et al. 2019). Still, male mating strategy could shift with size from courtship behaviors, where claw length may be more important, to coercion as males grow, which is supported by behavioral differences in courtship in Hughes (2011) and would obscure a generalized influence of size and claw length on male reproductive success.

Coloration - Conspicuous coloration commonly signals male health and competitive ability (e.g., McGraw and Ardia 2003; Plasman et al. 2015). Males with bright colors or specific hues experience increased fitness through female choice (e.g., Safran et al. 2005). Female color also may be important for male mate choice and reproductive success (Lüdtke and Foerster 2019). Indeed, color and health are associated in pond turtles of both sexes. For example, female red-eared slider turtles displayed decreased brightness of chin stripes following an immune challenge (Ibáñez et al. 2014). In painted turtles, stress biomarkers and immune function predict brightness and hue of forelimb stripes in sex- and size-dependent contexts (Judson et al. In review). Even so, we detected no substantive covariances between either male or female reproductive success and forelimb stripe color, which casts doubt on the long-presumed function of forelimb stripe coloration in pond turtles as a mate attractant (e.g., Ibáñez et al. 2014; Steffen et al. 2015; Judson et al. In review).

Forelimb stripe color might affect fitness separate from signaling mate quality in emydid turtles. Color may be a species recognition signal, such that heterospecific matings are reduced in areas where multiple sympatric species of similarly sized turtles interact (e.g., Vogt 1993), as is the case for much of the painted turtle’s geographic range (Ernst and Lovich 2009). Alternatively, painted turtle limb and head stripes may function in crypsis (Rowe et al. 2014), though no evidence of their cryptic advantage exists to date. Our methods precluded turtle visual system modeling (i.e., visual-system specific quantum catches) due to absence of UV measures, and thus we measured the forelimb stripes, which show little UV reflectance (Steffen et al. 2015) and predict health state in painted turtles (Judson et al. In review), so our results should not be limited by the absence of UV measures. However, head stripes, which we did not measure in this experiment and which have much greater UV reflectance (Steffen et al. 2015), might affect reproductive success in painted turtles. The impact of head stripe coloration on reproductive success should be assessed to further elucidate the role of coloration in mate choice of freshwater turtles.

Though color of forelimb stripes does not appear to influence reproductive success, we found support for the CORT-Fitness Hypothesis (Bonier et al. 2009) in female painted turtles. Females with higher baseline CORT concentrations laid fewer eggs and thus had lower fitness (Fig. 3). Importantly, this finding is based on measuring CORT after the nesting season, rather than right after a nesting event. Thus, in an otherwise aquatic turtle, our measures are distinct from immediate stress responses to terrestrial reproductive effort (e.g., Polich 2018) and may better reflect baseline levels of stress. Experimentally increased baseline CORT concentrations are associated with decreased female reproductive success in other reptiles, including garter snakes (Robert et al. 2009) and eastern fence lizards (MacLeod et al. 2018). Decreased offspring survivorship after application of CORT to recently oviposited painted turtle eggs suggests increased maternal CORT may also limit offspring fitness (Polich et al. 2018), further decreasing lifetime fitness of these iteroparous turtles. Interestingly, although baseline CORT and stripe brightness are negatively associated in these painted turtles (Judson et al. In review), stripe brightness was not associated with reproductive success. CORT is often proposed to be a mediator of signal honesty through its influence on allocation of resources toward self-maintenance and away from reproduction (e.g., ornamentation to attract mates, reviewed in Leary and Baugh 2020). Thus, CORT might affect female brightness by advertising reproductive quality to male painted turtles while also directly mediating maternal allocation to reproductive bouts.

Adult sex ratio - Environmental contexts strongly influence reproductive success in many species, and glucocorticoids might mediate the interaction of environmental stress with reproductive effort (Bonier et al. 2009). Environmental stressors, including lack of resources, extreme temperatures, and adverse social contexts, can affect CORT concentrations and reduce reproductive success (Henderson et al. 2017; Lea et al. 2018). Skewed ASRs can be another environmental stressor via increased mate competition or harassment from conspecifics (e.g., Le Galliard et al. 2005; Lea et al. 2018). We manipulated ASR of three experimental ponds to explore the influence of ASR on reproductive success of both sexes. We found no evidence that ASR affected female clutch size using a model selection approach, but we could not model factors influencing whether or not a female oviposited as only three females recovered from the ponds did not oviposit. However, two of these turtles exhibited some of the highest CORT concentrations of females in this study (Fig. 3), all three were kept in the female-biased ASR pond, in addition to one female that did not reproduce and was not recovered from the pond (Fig. 2), and turtles from this pond tended to have higher CORT concentrations compared to those from the equal and male-biased ponds (Fig. S5). Although not statistically significant, the effects of a female-biased ASR should be studied, particularly in this species and other turtles that have TSD. Climate warming presumably will produce increasingly female-biased ASR in such turtles, as warmer incubation conditions yield female hatchlings (Janzen 1994; Schwanz et al. 2010). The female-biased pond also did not yield any clutches with multiple paternity, which may suggest a lack of re-mating opportunities due to limited male availability (Uller and Olsson 2008).

We detected weak evidence that ASR influenced male reproductive success, as model weights were low. Males in the male-biased ASR pond achieved lower reproductive success on average than males in the female-biased pond (Fig. 4), and the variance in male reproductive success was greater in the female-biased pond than in the male-biased or equal ratio ponds (Fig. 2). This result contrasts with the hypothesis that increased male availability should enhance female choice and thus increase variance in male reproductive success (Kvarnemo and Ahnesjö 2002). A simulation study of male painted turtle reproductive success in varying ASR found that, consistent with our results, males in a female-biased population exhibited increased average reproductive success and increased variance in reproductive success (Hughes 2011). This pattern was attributed to increased male encounters of females in a female-biased population, allowing more opportunities for mating. Although female choice cannot be ruled out as a factor by us or by Hughes (2011), it is not necessary to invoke female choice as driving any of the reproductive patterns we detected with ASR treatments or with the phenotypes measured. Importantly, we cannot disentangle density effects from ASR effects, as there were differing numbers of turtles in each pond. Thus, future studies should evaluate whether density or ASR plays a greater role in reproductive success (Wacker et al. 2013), and whether females’ responses to prolonged skew in ASR include changes in use of stored sperm in future reproductive bouts.

We assessed relationships between multiple traits, ASR, and reproductive success in sexually dimorphic painted turtles under semi-natural conditions to understand how phenotypes and ASR might affect male and female reproductive success. We found strong evidence that female body size and CORT concentration influenced clutch size, whereas relationships for males were weaker and only suggested a trend toward male-biased ASR reducing male reproductive success. Despite many hypothesized relationships between the phenotypes quantified in this study (e.g., male claw length and forelimb coloration) and potential female mate choice in painted turtles, we detected no evidence of female mate choice. Thus, reproductive dynamics in turtles may be more complex than is often assumed.

## Acknowledgements

We thank R. Polich, T. Mitchell, D. Warner, N. Howell and J. Braland for pond building and maintenance at the ISU Horticulture Farm, B. Bodensteiner for camera equipment, C. Adams for nest monitoring, A. Toth, K. Roe, and J. Nason for project guidance and comments, R. Waples for guidance on Opportunity for Selection indices, E. Gangloff and A. McCombs for statistical advice, A. Bronikowski for manuscript feedback, and the many past and present members of the Janzen lab for the collection and care of turtles.

