## Supplemental Figures for "Phenotypic and environmental predictors of reproductive success in painted turtles"

Supplementary Material

Supplementary Table 1: Female model selection results ranked according to $AIC_c$
File name: FemaleModels.csv

Supplementary Table 2: Male model selection results ranked according to $AIC_c$
File name: MaleModels.csv

Supplementary Table 3: Male model selection results including only an intercept-only zero-inflation model ranked according to $AIC_c$
File name: MaleModelsConditionalOnly.csv

Supplementary Fig. 1: Model fit test for full female model

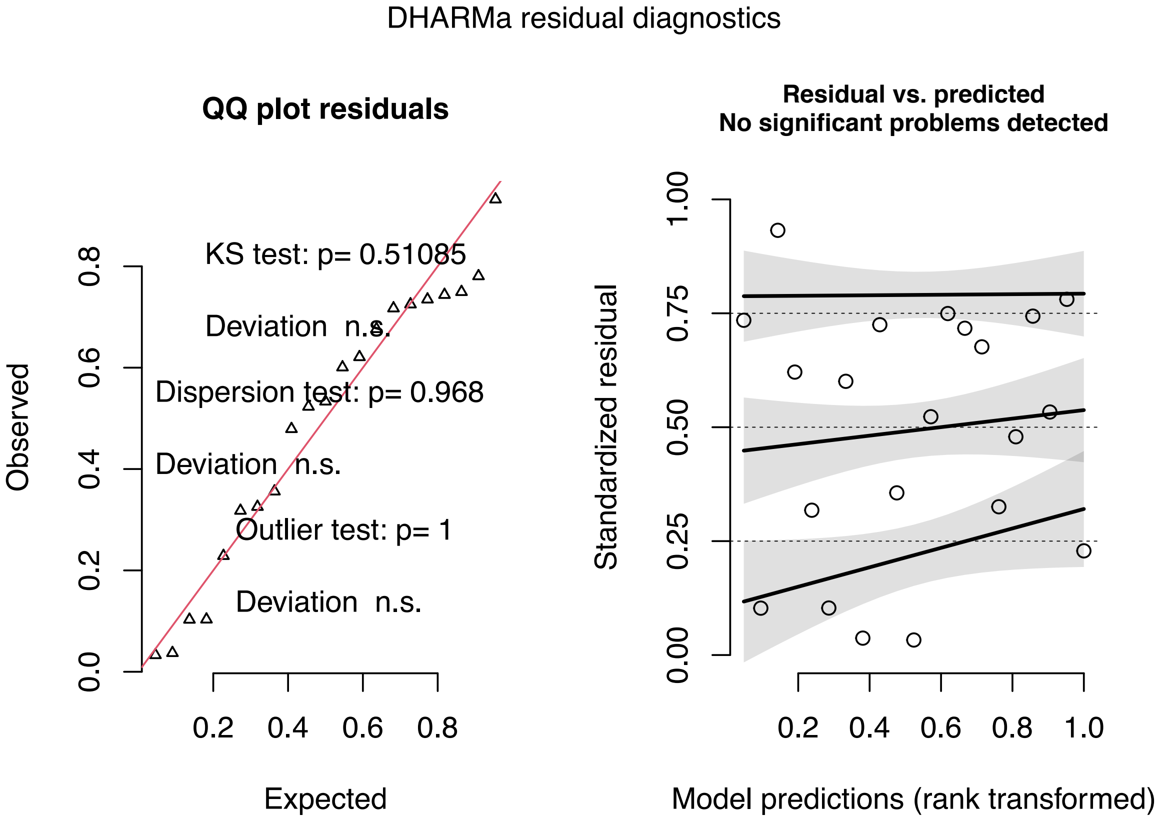

Supplementary Fig. 2: Model fit test for best-fitting female model

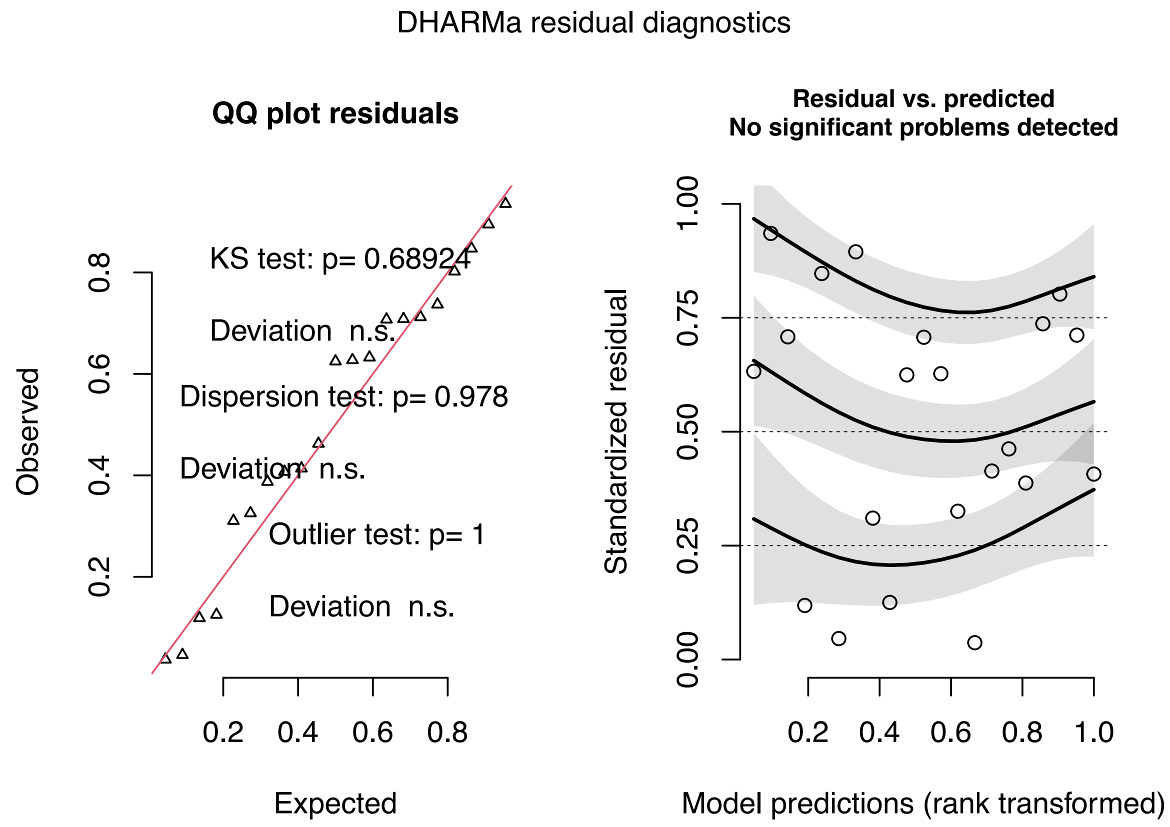

Supplementary Fig. 3: Model fit test for full male model

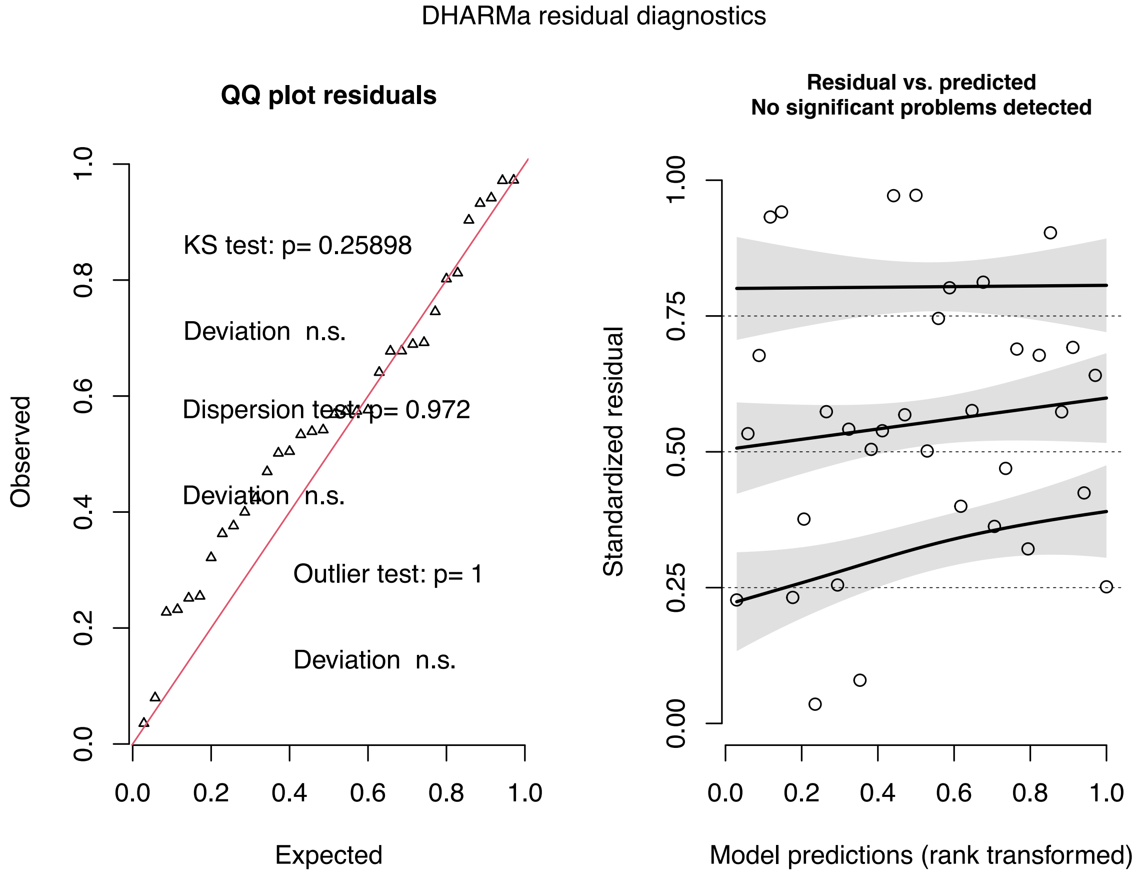

Supplementary Fig. 4: Model fit test for best-fitting male model

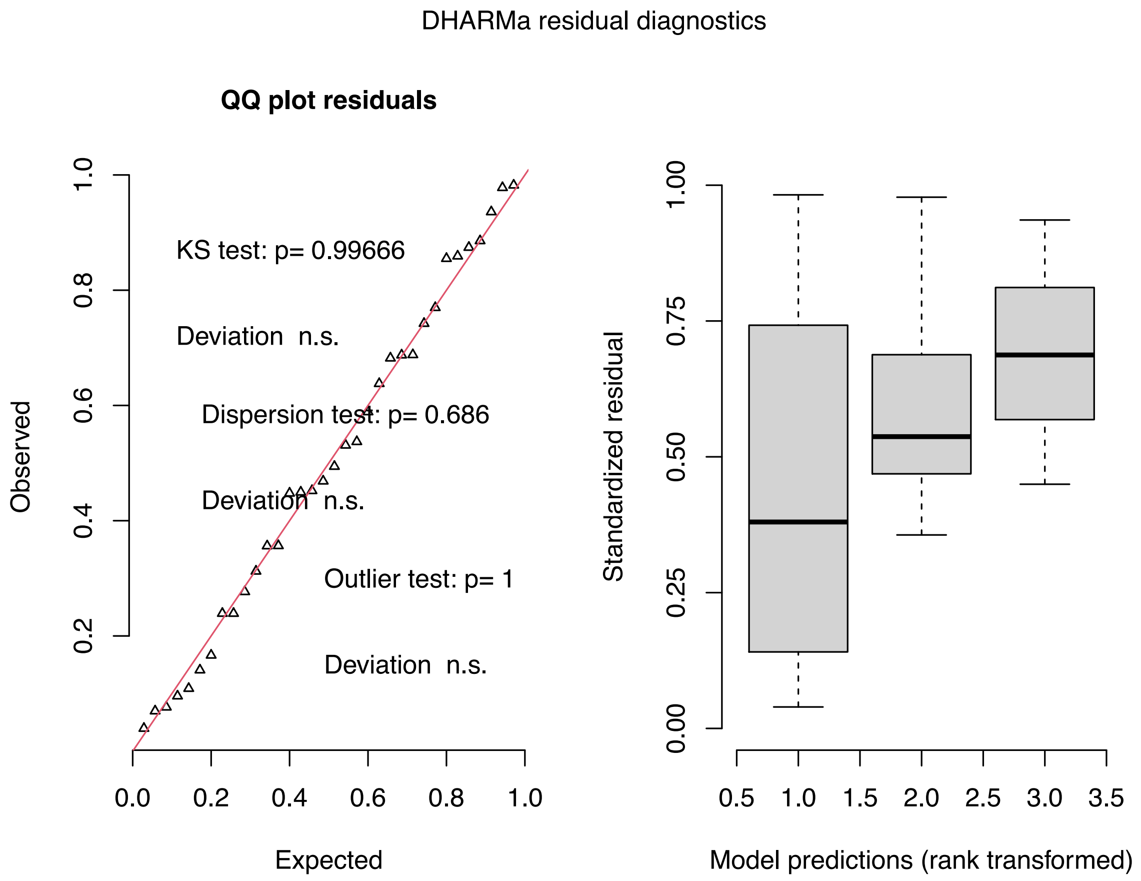

Supplementary Fig. 5: Comparison of female baseline corticosterone (CORT) concentrations across adult sex ratio (ASR) treatments.

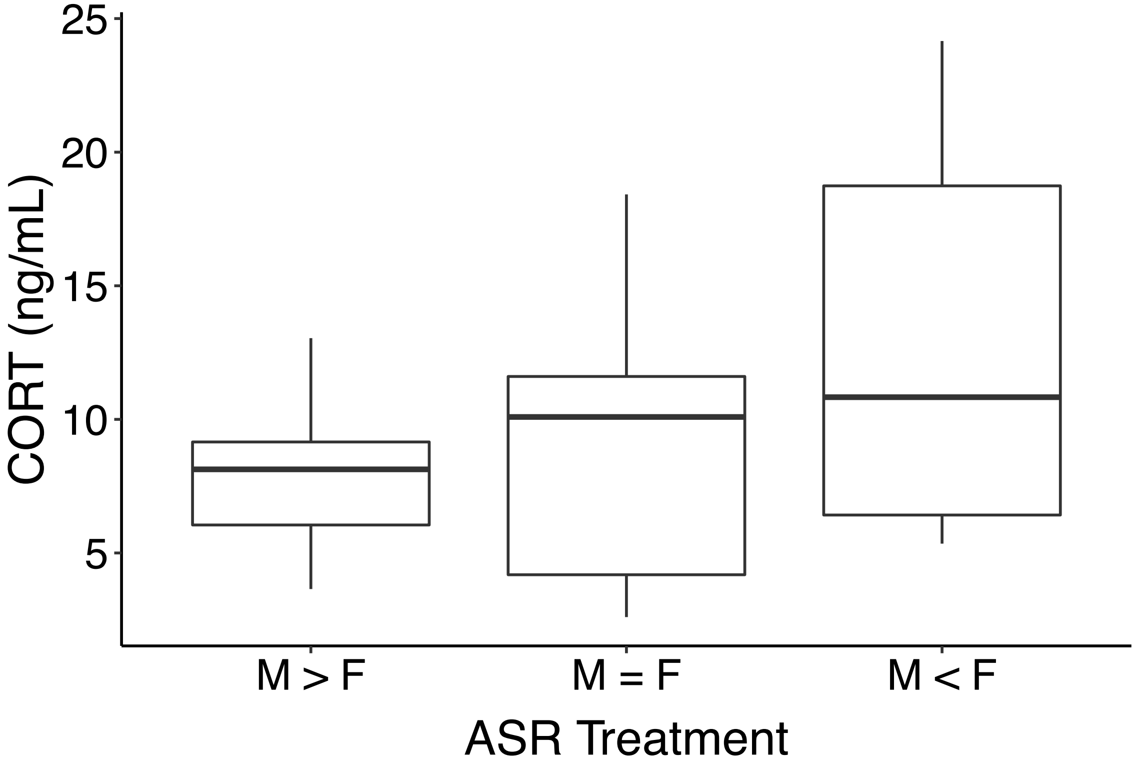
